# Myosoft: an automated muscle histology analysis tool using machine learning algorithm utilizing FIJI/ImageJ software

**DOI:** 10.1101/703736

**Authors:** Lucas Encarnacion-Rivera, Steven Foltz, H. Criss Hartzell, Hyo-Jung Choo

**Author notes:** Corresponding author, Hyo-Jung Choo, Ph.D.: Emory University, Department of Cell Biology, 615 Michael street, Rm 542, Atlanta, GA 30322, USA.

## Abstract

**Background:** Skeletal muscle is comprised of a heterogeneous population of muscle fibers which can be classified by their metabolic and contractile properties (fiber “types”). Fiber type is a primary determinant of muscle function along with fiber size (cross-sectional area). The fiber type composition of a muscle responds to physiological changes like exercise and aging and is often altered in disease states. Thus, analysis of fiber size and type in histological muscle preparations is a useful method for quantifying key indicators of muscle function and for measuring responses to a variety of stimuli or stressors. These analyses are near-ubiquitous in the fields of muscle physiology and myopathy, but are most commonly performed manually, which is highly labor- and time-intensive. To offset this obstacle, we developed Myosoft, a novel method to automate morphometric and fiber type analysis in muscle sections stained with fluorescent antibodies.

**Methods:** Muscle sections were stained for cell membrane (laminin) and myofiber type (myosin heavy chain isoforms). Myosoft, running in the open access software platform FIJI (ImageJ), was used to analyze myofiber size and type in transverse sections of entire gastrocnemius/soleus muscles.

**Results:** Myosoft provides accurate analysis of muscle histology >50-times faster than manual analysis. We demonstrate that Myosoft is capable of handling high-content images even when image or staining quality is suboptimal, which is a marked improvement over currently available, comparable programs.

**Conclusions:** Myosoft is a reliable, accurate, high-throughput, and convenient tool to analyze high-content muscle histology. Myosoft is freely available to download from Github at https://github.com/Hyojung-Choo/Myosoft/tree/Myosoft-hub.

## Introduction

Skeletal muscle is the most abundant tissue and is responsible for movement and posture [1, 2]. The human musculature comprises over 600 skeletal muscles, which generate a diverse range of contractile forces due to differences in the compositions of their constituent muscle fibers [3, 4]. Muscle fibers are broadly classified by contractile kinetics (slow or fast twitch, referred to as type I or type II, respectively). Type II fibers may be further categorized according to metabolic activity (oxidative or glycolytic) [5, 6]. According to this method of classification, there are four general fiber types: type I fibers (slow twitch/oxidative metabolism), type IIa (fast twitch/oxidative metabolism), and types IIx and IIb (faster and fastest twitch/glycolytic metabolism) [5, 7]. Each fiber type contains different myosin heavy chain (MyHC) isoforms, which differ with respect to ATPase activity and contraction speed. The contractile properties of a muscle fiber can be inferred by its size (generally reported as its cross-sectional area, CSA) and type [4, 5]. The sizes and types of fibers in a given muscle collectively contribute to its functional output [8–10].

The distributions of fiber size and type display plasticity in response to physiological pressures like aging and exercise and are altered in cases of neuromuscular disease [11–15]. Thus, analysis of fiber size and type in histological muscle preparations can be a useful method for quantifying key indicators of muscle function and for measuring responses to a variety of stimuli or stressors. However, despite the value of such analysis, it is often performed manually, which is both labor-intensive and time-consuming. To offset this obstacle, several groups have developed software that automates analysis of muscle histology [16–19]. Unfortunately, these methods may incur a significant learning curve for the investigator. Additionally, all current methods consist of a single round of image segmentation and depend on performing sequential transformations on the original image. This puts greater emphasis on the quality of the stain and appearance of the image to achieve accurate analysis. Machine learning offers unprecedented potential in resolving these present limitations in automated image analysis [20]. Recently, machine learning based tools for image segmentation, like the Trainable Weka Segmentation (TWS) tool, have been developed, offering a novel avenue for image analysis. A TWS classifier can be trained to recognize and distinguish between the muscle fiber boundary and intra-fiber space after a few simple manual annotations that instantiate these two features. Then, given an image, the classifier predicts the muscle fiber boundary and intra-fiber space. The results of this prediction are represented as a probability rendering of the original image, where darker pixels represent higher probabilities for the boundary, and lighter pixels represent lower probabilities. Since the classifier is trained specifically to segment images of muscle tissue, it can “learn” to account for and clarify flaws in an immunostained image that would otherwise confound the analysis or necessitate additional segmentation methods.

Here, we present Myosoft, a novel tool to analyze muscle histology that synergizes machine learning-based image segmentation with thresholding-based object extraction and quantification. Myosoft uses pre-trained machine learning classifiers to delineate muscle fiber boundaries and subsequently extracts the size, type, and relevant morphometric features of the fiber. Additionally, Myosoft is run in the open-access image analysis software FIJI (Fiji is Just ImageJ) which is widely used to analyze cellular histology [21, 22]. Altogether, Myosoft is a high-throughput, quick, accurate, and convenient solution to analyzing large sections of muscle tissue, capable of circumventing the error, bias, and labor incurred by manual annotation.

## Methods

### Mice and muscle tissue preparation

All experiments involving animals were performed in accordance with approved guidelines and ethical approval from Emory University’s Institutional Animal Care and Use Committee. C57BL/6 and *Dmd<mdx-4Cv>* mice were purchased from Jackson Laboratories. Six-month-old male mice were used for all experiments. Mice were euthanized via inhalation overdose of isoflurane, the skin was removed from the hindlimbs, and gastrocnemius muscles were excised. Gastrocnemius muscle tissues were mounted in OCT freezing medium (Triangle Biomedical Sciences), snap-frozen in liquid N2-cooled 2-methylbutane and stored at −80°C for cryo-sectioning. Tissue cross sections of 10 μm thickness were collected every 400 μm using a Leica CM1850 cryostat.

### Immunofluorescent staining

For immunostaining specific types of myosin heavy chain and laminin, tissue sections were first treated with mouse-on-mouse reagents (M.O.M. Kit, Vector Laboratories Inc.) to block endogenous Fc receptor binding sites followed by a 1 hour incubation with 5% goat serum, 5% donkey serum, 0.5% BSA, 0. 25% Triton-X 100 in PBS (blocking buffer). Sections were then labeled with an undiluted 1:1:1 mixture of mouse monoclonal antibodies BA-D5 (anti-MYH7, fiber type I), SC-71 (anti-MYH2, fiber type IIa), and BF-F3 (anti-MYH4, fiber type IIb) (hybridoma supernates, Developmental Studies Hybridoma Bank) supplemented with rabbit polyclonal anti-laminin antibody (2 μg/ml, Sigma) overnight at 4°C. Control sections were incubated with species-matched non-immune IgGs. Sections were then incubated with isotype-specific Alexa Fluor (AF) conjugated secondary antibodies: anti-mouse IgG2b-AF647, anti-mouse IgG1-AF488, anti-mouse IgM-AF350, and anti-rabbit IgG-AF594 (Invitrogen, Molecular Probes) to mark type I, IIa, IIb fibers and laminin, respectively. Sections were mounted using ProLong Diamond anti-fade mountant (ThermoFisher Scientific).

### Image acquisition

All images were obtained using a Nikon Eclipse Ti-E inverted epifluorescent microscope equipped with a motorized stage. Images were acquired in NIS-Elements software (Nikon) with a 10x/0.3NA PlanFluor objective. The ND acquisition menu within Elements was used to take images from adjacent fields of view and digitally stitch them (with 15% overlap) to form a single image of the entire muscle crosssection (approximately 20 – 30 mm^2^) used for analysis.

### Image analysis

#### Manual Outline and Fiber Typing

Four images containing 150-200 fibers were taken from areas of the muscle section where all fiber types were represented. CSA measurement (using the polygon tool) and fibertyping (using the counting tool) was performed for all fibers in the images (excluding those on the edge) by two individuals in Fiji.

#### Images used to test the efficacy of Myosoft and compare it to other programs

Images for comparison of analysis programs were generated by fractionating large, whole tissue section images. Small images (containing 200-1000 fibers) were chosen after dividing the original image into sixteenths while larger images (1000+ fibers), were taken from 2 different muscle sections divided into halves or fourths. Selection was accomplished with a random number generator to eliminate bias from the process. If an image was randomly selected and had significant fluorescence artifacts or tissue damage, this area was either excluded, or a new image was chosen. Each of these images was then run through Myosoft and its peer programs to obtain simple fiber counts. False negatives and false positives were then manually scored for each image. All above images were then classified as either poor or good quality. Stain quality was quantified as the ratio of intensity between the intra-fiber space and fiber boundary. Ratios >5 were defined as good stain quality while ratios <5 were defined as poor stain quality. Measurements were made at various locations to account for non-uniformity in staining within single images.

### Statistics

An unpaired Student’s *t*-test was used to determine the statistical significance between two groups. The significance of differences between multiple groups was evaluated by one-way ANOVA with Bonferroni’s post-test correction. Fiber size distributions were analysed using Kruskal-Wallis non-parametric ANOVA. Histogram bin sizes were determined via the Freedman-Diaconis rule: 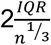, where IQR is the interquartile range and n is the total number of observations (fibers) taken from WT gastrocnemius/soleus [23]. All statistical comparisons were performed using Prism 7 software (GraphPad Software, Inc). A p-value of <0.05 was considered significant.

### Myosoft download and tutorial

The code for Myosoft (an ImageJ macro) is freely available and can be accessed directly in Supplemental file 1 along with a tutorial and troubleshooting instructions (Supplemental File 2), or downloaded through the Choo lab repository on GitHub (https://github.com/Hyojung-Choo/Myosoft/tree/Myosoft-hub).

## Results

### Four components of the Myosoft pipeline

The Myosoft image analysis pipeline features 4 distinct modules: image pre-processing, segmentation, thresholding, and region of interest (ROI) overlay. In the preprocessing step, simple contrast enhancement is applied to the membrane-stained channel of the image and a 4×4 convolutional matrix is applied to enhance the cell boundary edges (Fig. 1 Step 1 and 2). Next, to facilitate computation on computers with limited CPU clock speed and RAM, the entire 10x muscle image is sliced into 4 to 25 (that is, 2^2^ to 5^2^; we use 4^2^ = 16) equally sized smaller images that are then processed independently (Fig. 1 Step 3). Although not strictly necessary for the analysis presented here, this step drastically reduces the computing power required for execution of the Myosoft macro and is thus highly recommended (and a default setting in Myosoft). In the segmentation step, sliced images from step 3 are segmented using a pre-trained machine learning classifier (primary classifier). The classifier predicts fiber boundary and intra-fiber space using pixel intensity values from the laminin membrane stain. Each of the slices are segmented and saved (Fig. 1 Step 4). Next, images are subjected to another round of segmentation using a separate machine learning classifier (iterative classifier). The iterative classifier was trained on images already segmented by the primary classifier. The output of this step is an image with defined and continuous fiber boundaries and intra-fiber spaces with minimal noise (Fig. 1 Step 5). In the thresholding step, Myosoft retrieves all iteratively segmented images and stitches them to re-form a unified and complete segmented image of the original muscle section (Fig. 1 Step 6). Next, a maximum entropy thresholding algorithm is applied to this image to binarize it (Fig. 1 Step 7). In the binary image, pixels are either black (representing the membrane-stained cell boundary) or white (corresponding to cytoplasm of myofibers, which are unstained). A particle analyzer extracts contiguous white pixels as objects and represents them as ROIs (Fig. 1 Step 8, red). The ROIs obtained after gating in step 8 are expanded according to an adjustable ROI expansion factor to closely match the fiber boundary marked by laminin in the input image (Fig. 1 Step 8, yellow). In the final ROI overlay step, original MyHC channel images are retrieved and expanded ROIs are overlaid. Myosoft then extracts several measurements within each ROI from each channel, including mean intensity, mode, and standard deviation. The numerical data for each channel image is saved as an XLS (Microsoft Excel format) file. Additionally, a reference image for each channel is saved which illustrates all the ROIs and their ID number (Fig 1. Step 9). Each ROI has its own numerical ID which is indexed identically between the data and the reference images. ROIs of whole muscle sections are color-coded according to CSA and the color-coded image is saved (Fig. 1 Step 10). The distribution of intensity values on each channel is plotted by the user to sort fiber types and to obtain CSA information. (Fig 1. Steps 11 and 12)

**Figure 1.**
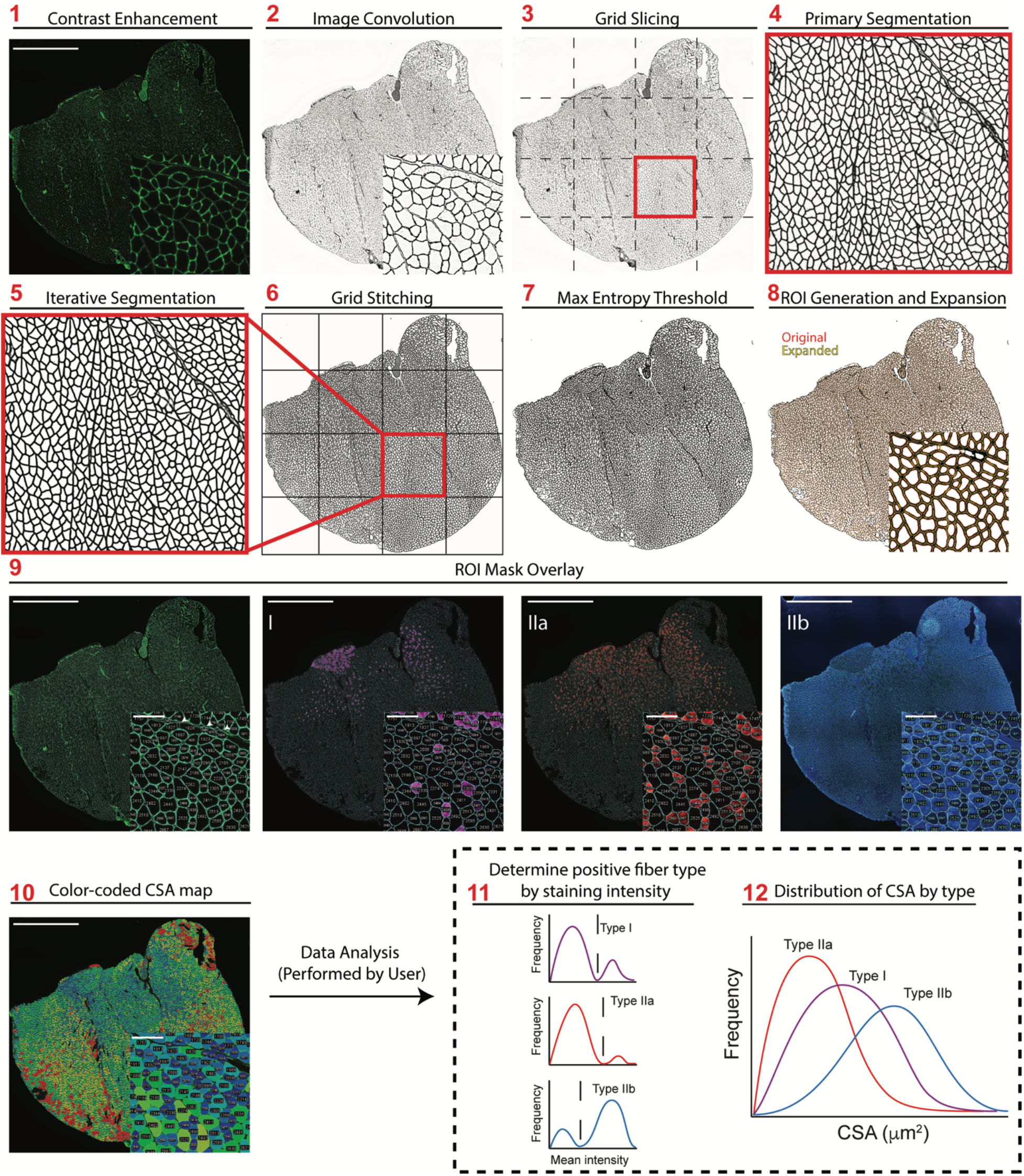
Overview of Myosoft Image Analysis Pipeline. **Step 1** Image of laminin stain with contrast enhancement performed. **Step 2** Laminin stain following 8-bit conversion and image convolution. **Step 3** Image is cut in a 4×4 grid to create 16 equally sized images. **Step 4** Probability map following application of the primary machine learning classifier. **Step 5** Probability map following application of the iterative machine learning classifier. **Step 6** Recombination of all iteratively segmented grid images to form segmented full-tissue image. **Step 7** Segmented image following pixel binarization using max entropy threshold. **Step 8** Initial ROI mask (red) acquired using particle analysis and mature ROI mask (yellow) following ROI enlargement. **Step 9** Original single channel images with indexed ROI overlay. **Step 10** Laminin stain overlaid with ROI heat map color coded by size. **Step 11** Intensity histograms (generated by user) identify intensity thresholds for each fiber type. **Step 12** CSA distributions for each fiber type (generated by user), made possible through Myosoft ROI indexing. Scale bars: 1500μm, large images; 100μm, inset images.

### Adjustable Parameters

In our initial tests of Myosoft, we noticed that some objects that were not myofibers were erroneously scored as myofibers (see white arrowheads in left panel of Fig. 1 Step 9). However, by visual inspection, it is clear that these objects differ in size and shape from true muscle fibers. We reasoned that exclusion criteria based on morphometric features of objects would eliminate these artefacts from detection. Therefore, Myosoft prompts the user to enter constraining values for several parameters that correspond to specific morphometric features. This is accomplished through the Extended Particle Analyzer plugin within the Biovoxxel Toolbox. Parameters include cross-sectional area (Fig. 2A), solidity (which measures the ratio of an object’s area to that of a convex polygon of equivalent height and width, Fig. 2B), and circularity (a measure of how well an object approximates a circle, Fig. 2C). Adjusting these parameters can be used to exclude artifacts (e.g., interstitial spaces), objects that are not fibers (e.g. blood vessels), and improperly annotated fibers (e.g., two fibers where the boundary between them is incomplete, Fig. 2D).

**Figure 2.**
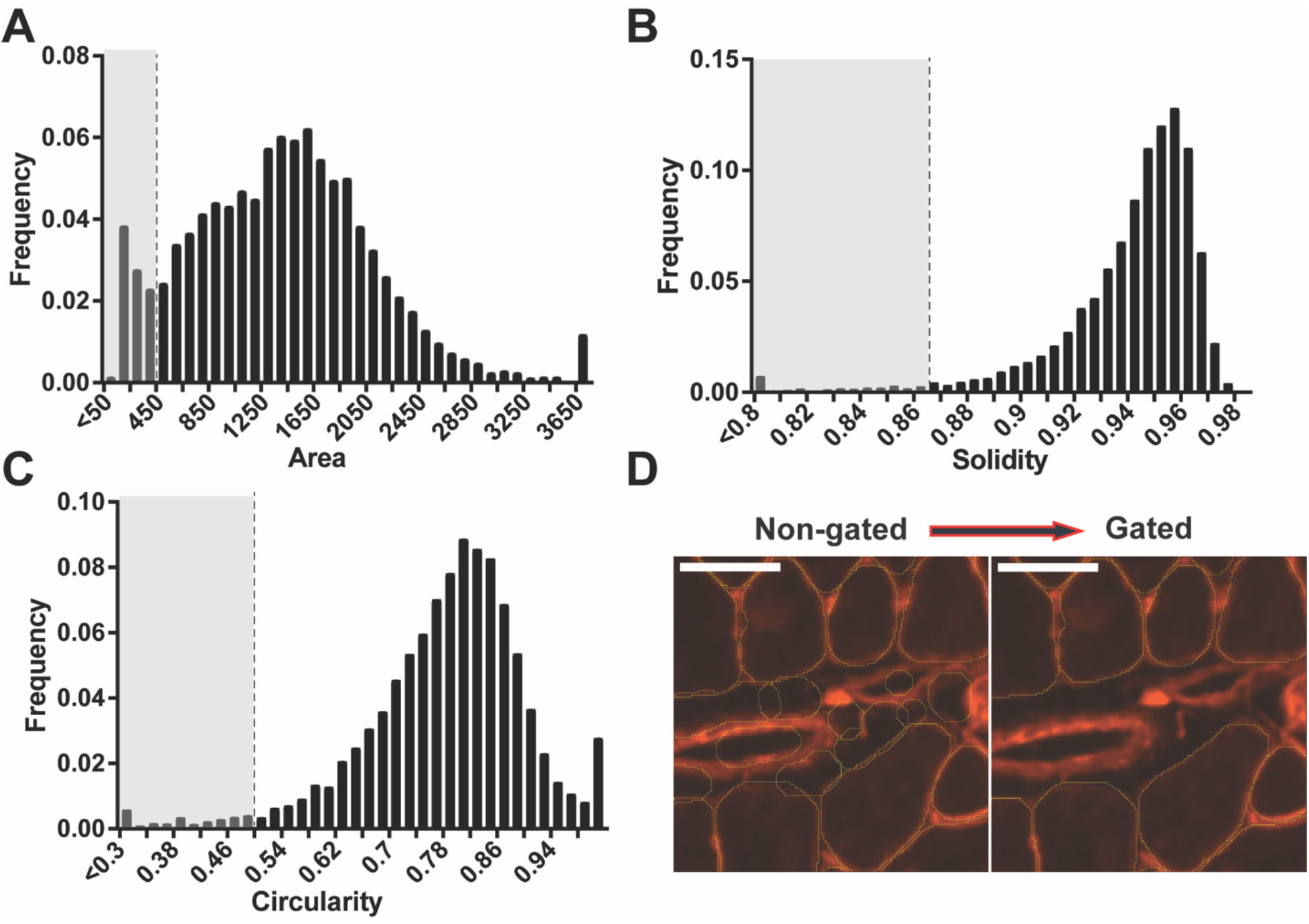
Morphometric gates used to exclude false positives. **a** Frequency distribution of area when no morphometric gates were used with an arbitrary threshold equal to the default area-gate minimum. **b** The frequency distribution of solidity when no morphometric gates were used with an arbitrary threshold equal to the default solidity-gate minimum. **c** The frequency distribution of circularity when no morphometric gates were used with an arbitrary threshold equal to the default circularity-gate minimum. **d** ROI mask when no morphometric gating is used vs. when it is used (scale bars = 50μm).

### Fiber typing is determined by gating of MyHC fluorescent intensity distributions

We stained muscle fiber types I, IIa, and IIb using specific MyHC antibodies for each type (Fig. 3A-C). As mentioned above, Myosoft will return several measurements for each ROI on each color channel. To perform fiber type analysis, we first plotted a frequency distribution of intensity values for each channel (Fig 3A-C). The distributions were typically bimodal, with peaks corresponding to myofibers that are positive or negative for each MyHC isoform. From the intensity histograms, we established thresholds that were used to define positive and negative fibers of each type (Fig. 3A-C, dotted vertical line).

**Figure 3.**
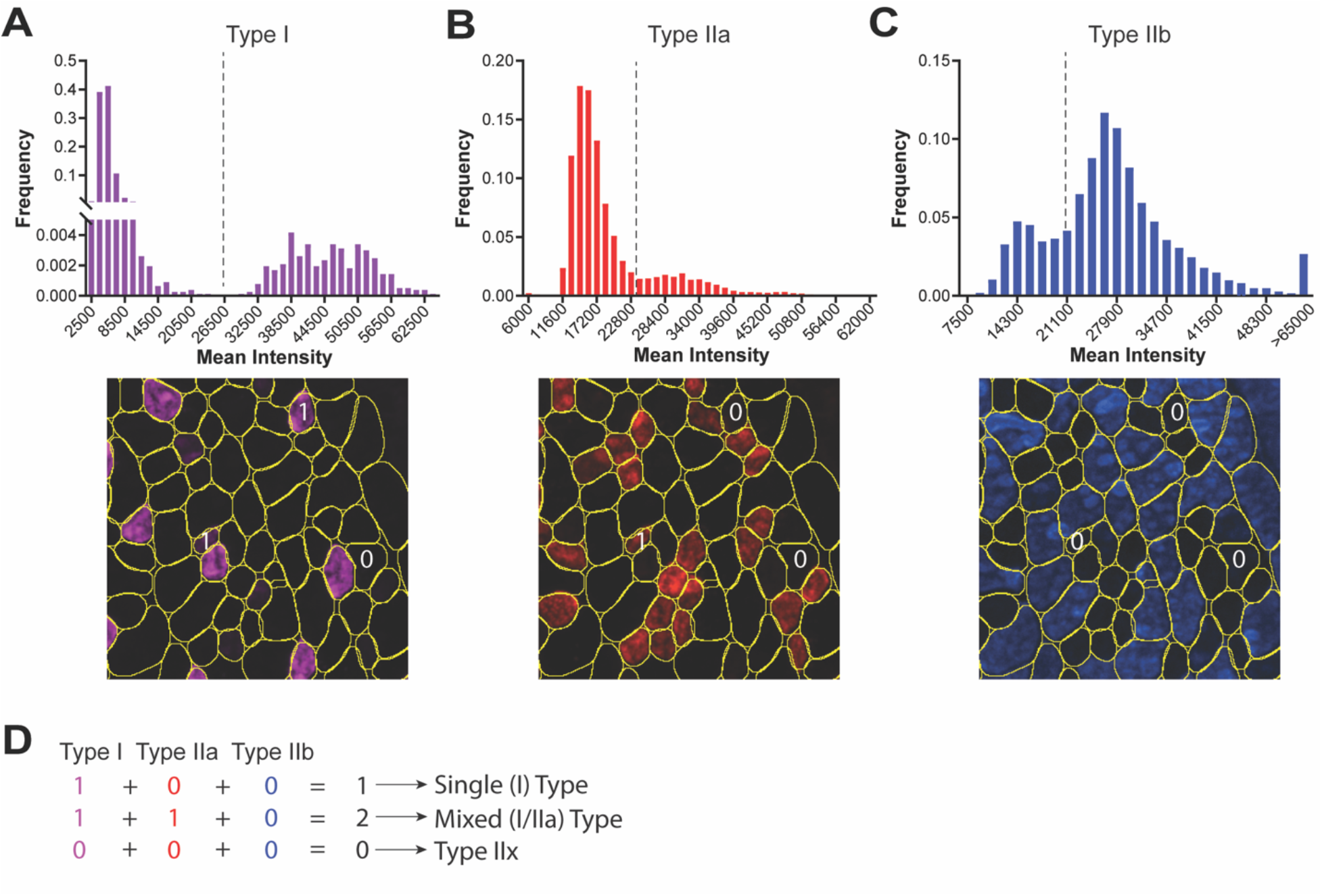
Method for extracting fiber type data from Myosoft measurements. Mean intensity histograms for type I (**a**), type IIa (**b**), and type IIb (**c**) show bimodal distributions, with peaks corresponding to low and high mean intensity values. A threshold is applied (vertical dotted line), and all values above the threshold are considered positive for that fiber type. **d** Logic for identifying type IIx fibers. Fibers that are marked as positive on any channel receive a value of 1 for that channel while fibers marked as negative receive a value of 0. Type IIx fibers are those fibers that receive a score of 0 for all 3 channels (sum=0, bottom), while mixed fiber types are indicated by sums>1 (middle).

Because our images have only 3 color channels, no antibody was used to label type IIx fibers; instead, the presence of type IIx fibers was inferred through the absence of any other MyHC isoform staining. To identify IIx fibers, it was thus necessary to determine which fibers had sub-threshold intensity values for every channel. To that end, we generated a logical array from Type I, IIa, and IIb intensity measurements, assigning a value of 1 to intensity measurements above the threshold on a given channel and 0 to intensity measurements below the threshold on that channel (example fibers are indicated in Fig 3A-C). For all fibers, it is possible to create a new array equal to the sum of the other three arrays (Fig 3D). In this array, type IIx fibers will be those with a value of 0, since they were determined to be negative for all fiber types (Fig 3D). This also allows for the detection of mixed fiber types, which are represented as any fiber in the array with a value greater than 1 (Fig 3D).

### Myosoft Performance is comparable to manual analysis

To evaluate the accuracy of Myosoft, we assessed the comparative performance between manual analysis and Myosoft with respect to their ability to correctly identify muscle fibers. We segmented the original image of the muscle section into 16 equally sized images and randomly selected 4 of these images for analysis. False positives are defined as ROIs that do not delimit a single, whole muscle fiber (where “true” fibers are determined with manual annotation). We note three possibilities for this type of error: a single ROI outlining 2 fibers (2 as 1), 2 ROIs outlining a single fiber (1 as 2), and ROIs marking objects that are not fibers (nonfiber) (Fig. 4A). The false positive rate (0.35%) is thus the number of false positives divided by the total number of fibers counted via manual analysis. Conversely, the false negative rate is defined as the rate at which the program does not generate an ROI for a fiber (Fig. 4A, 0.95%).

**Figure 4.**
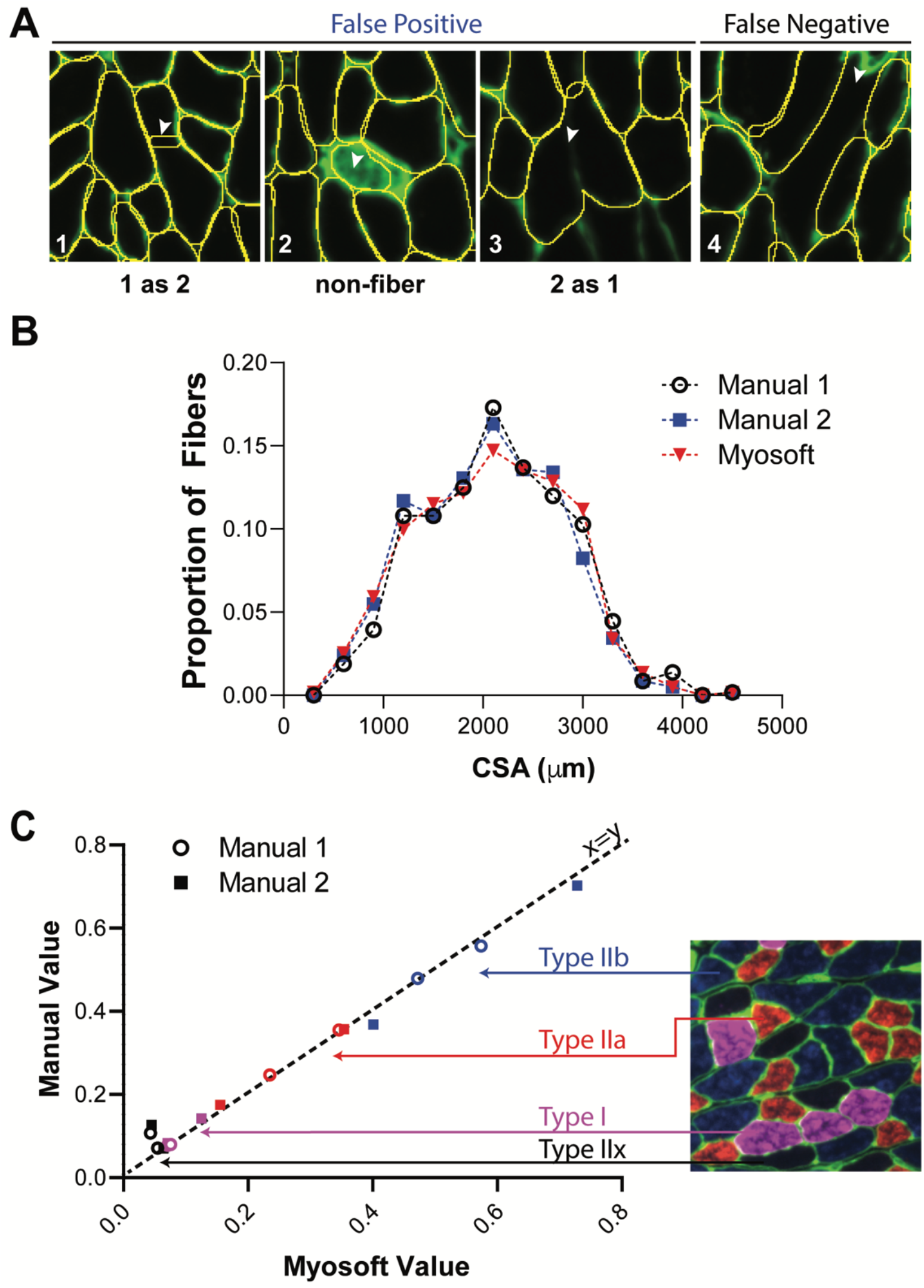
Myosoft is comparable to manual analysis. **a** Operational classifications of false positive and negative. **b** Myofiber CSA distributions were not different when determined with Myosoft or by manual annotation (p>0.9, Kruskal-Wallis non-parametric ANOVA with Dunn’s multiple comparison test). Results of manual analysis from 2 investigators are shown. **c** Proportion of each fiber type in a given muscle section determined manually or using Myosoft. Proportions determined manually are on the y-axis and proportions determined by Myosoft are on the x-axis. Type I fibers indicated by purple symbols, Type IIa indicated by red symbols, Type IIb indicated by blue symbols, and Type IIx indicated by black symbols.

We next wanted to determine how effective Myosoft is as a tool for automated myofiber size and type analysis. Two researchers with previous experience analyzing myofibers used the polygon tool in Fiji to outline muscle fibers from four 10X fields of view (~600 fibers total) to obtain CSA values. We then ran the same images through the Myosoft program and obtained a distribution of CSA across the images. The CSA distributions did not differ significantly between manual and Myosoft analysis (Fig. 4B). Next, we tested the accuracy of fiber typing using Myosoft. Fiber type analysis was manually performed on 4 images representing ~3000 fibers. We then used Myosoft to obtain mean intensity data for green, blue, and far red channels across these four images for fiber typing. The relative proportions of each fiber type was strongly correlated between Myosoft and manual analysis (r^2^ > 0.99) (Fig. 4C).

### Myosoft is a reliable program to analyze large-scale muscle histology images

Several tools exist for automation of muscle histological analysis, but Myosoft is the first we are aware of that employs machine learning. To validate this approach, we compared the performance of Myosoft and other programs that analyze muscle histology. We chose three recently published programs for initial comparison: SMASH (Smith and Barton 2014), Myosvision (Bergmeister, Groger et al. 2016), and MuscleJ

(Mayeuf-Louchart, Hardy et al. 2018). First, we compared the muscle fiber count across programs with manual count, which is considered here to be the “true” muscle fiber count. We chose 6 images, 4 of which were the randomly chosen images used to obtain the false positive and false negative rates, and the remaining 2 of which were images from 2 other muscle sections. While all muscle histology programs performed well counting myofiber number from muscle sections containing less than 500 fibers, Myovision identified fewer fibers from images containing ~750-1500 fibers, and Myovision, SMASH and Muscle J were all inaccurate in counting from images containing > 2,000 fibers. Myosoft was the only program to consistently reflect manual count throughout images regardless of number of fibers (Fig. 5A).

**Figure 5.**
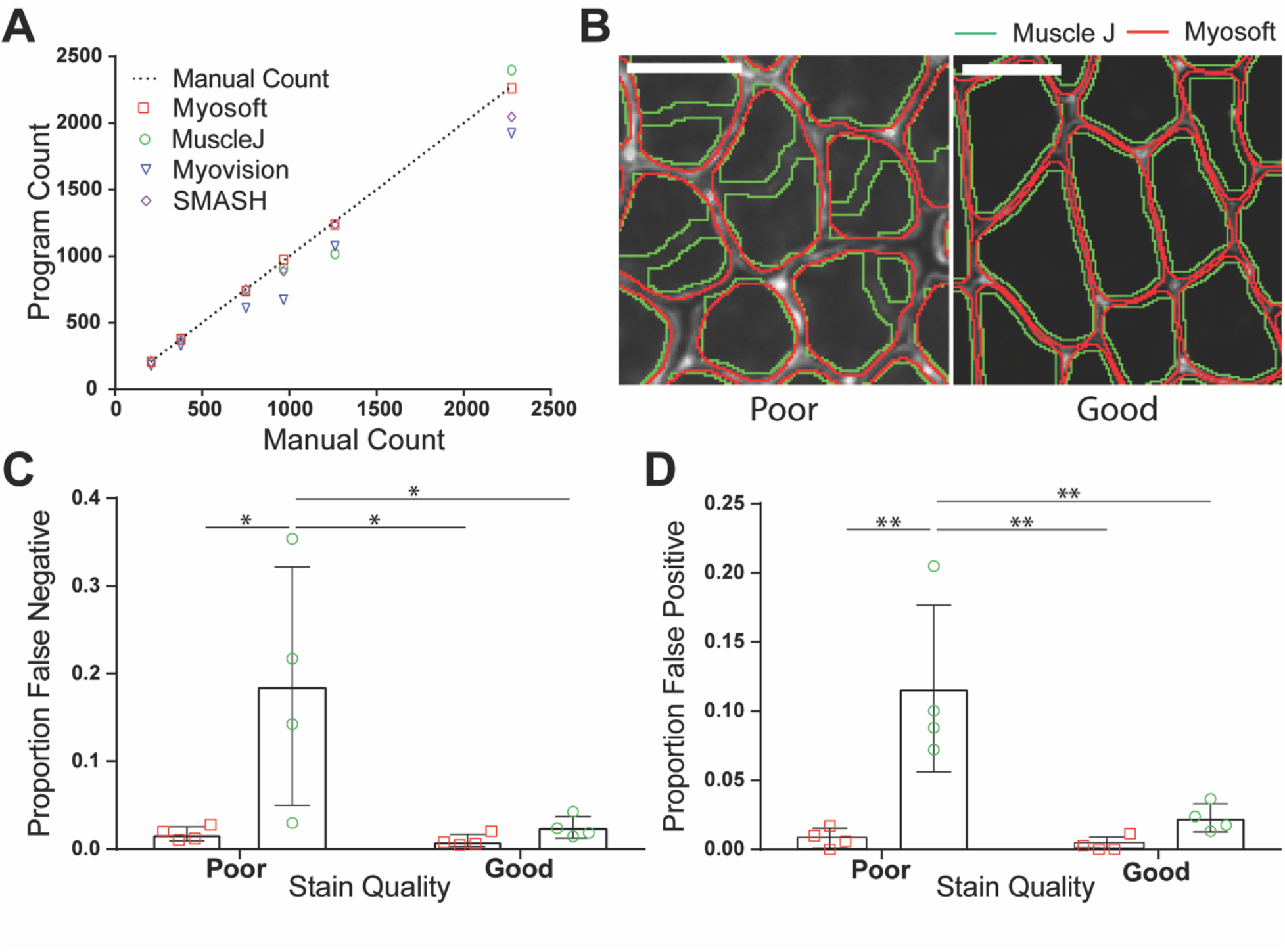
Efficacy of Myosoft compared to similar programs. **a** linear regression between manually counted fibers and the fiber count produced by SMASH, MuscleJ, Myovision, and Myosoft. **b** ROI outlines generated using Myosoft or MuscleJ in good-or poor-quality stains (scale bars = 50 μm). Good quality stains are defined as those stains which have >5-fold intensity relative to nearby un-stained space. **c** Proportion of false negatives generated between Myosoft and MuscleJ for good- and poor-quality stains. **d** Proportion of false positives generated between Myosoft and MuscleJ for good- and poor-quality stains.

Since these data only compare the raw fiber count with manual analysis, they do not represent the accuracy of the program *per se.* For example, a program may possess a high false positive and false negative rate, making the count appear artificially similar to the manual count but may not provide an accurate fiber identification. This problem may be more serious if the quality of the stain is not optimal. Since consistently producing high quality (i.e. high contrast) stains may be impractical, utilizing a program that can retain good performance on lower quality stains is desirable. Thus, we sought to examine how the false positive and false negative rate compares when the stain is “poor” and “good” between Myosoft and MuscleJ (Fig. 5B-D). We chose to compare Myosoft only to MuscleJ since it is the most recent muscle fiber analysis program and has the largest suite of abilities presented to date. Furthermore, it is like Myosoft in that it is capable of analyzing large-scale images, is coded in IJMacro and runs in Fiji (SMASH and myovision require a MATLAB compiler). We chose four images randomly from both “poor” and “good” quality stains. Good quality is defined as little to no noise between the muscle fiber boundary and intra-fiber space (the ratio of intensity between the intra-fiber space and fiber boundary > 5), while poor quality is defined as mid to high noise (intensity ratios < 5) between the muscle fiber boundary and intra-fiber space (Fig. 5B). When instances of false positive and false negative were counted manually for both Myosoft and MuscleJ, Myosoft displayed robust low false negative and false positive rates (<1.5 %) regardless of staining quality (Fig. 5C-D). Meanwhile, when the stain quality was poor, MuscleJ (Green line in Fig. 5B) performed significantly worse than Myosoft (Red line in Fig. 5B). With low quality images, MuscleJ’s false negative rate was ~20% and false positive rate was ~11%, which were significantly greater than those of Myosoft for both good- and poor-quality stains (Fig. 5C-D).

### Example: Myosoft clearly distinguishes normal and dystrophic muscle

Finally, we sought to evaluate the power of Myosoft to detect differences between normal and dystrophic tissue. For this purpose, we analyzed gastrocnemius sections from 20 week old WT or *mdx* mice. The *mdx* mouse is one of the most common muscular dystrophy models, harboring a spontaneous mutation in the *Dmd* gene (encoding dystrophin) and presenting with a moderate muscle phenotype [24]. *Mdx* muscle showed clear variability in its fiber size distribution, with abnormally high proportions of both atrophic and hypertrophic fibers. This feature, which is a hallmark of muscular dystrophy, was evident by inspection of the color-coded section image generated by Myosoft and through comparison of CSA histograms from *mdx* or WT muscle fibers (Fig. 6 A, C). When the distribution of fiber size was segmented according to type, a clear shift in *mdx* oxidative and glycolytic fiber size populations were observed. Type I and IIa fibers were, overall, still smaller in *mdx* muscles, but a subpopulation of these fibers was skewed to larger sizes. In contrast, types IIb and IIx fibers were conspicuously smaller in *mdx* muscle and more closely mimicked the general fiber size distribution (although these types also had minor subpopulations of very large fibers, with CSA >6000 μm^2^). Along with shifts in size, Myosoft also detected an alteration in the proportion of type IIa fibers, with *mdx* muscle containing about half the type IIa fibers of WT muscle.

**Figure 6.**
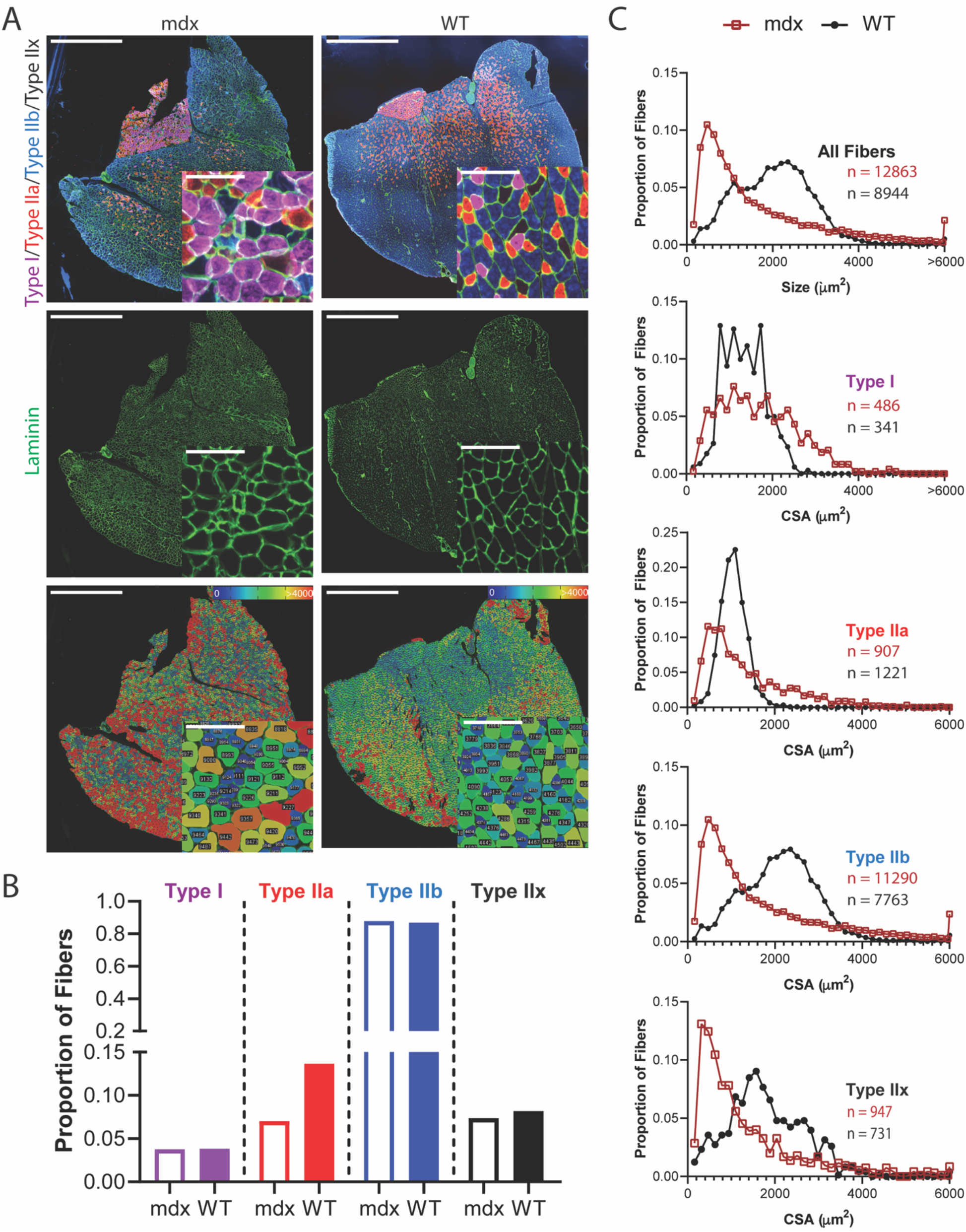
Myosoft is a convenient tool for detecting differences between control and dystrophic tissue. **a** Whole section images of dystrophic (*mdx*, left) or WT (right) gastrocnemius and soleus muscles showing fiber type distributions (top), laminin, marking cell boundaries (middle), or section maps color-coded according to fiber sizes (bottom). Scale bars: 1500μm, large images; 100μm, inset images. **b** Fiber type proportions determined by analysis of gastrocnemius/soleus sections from *mdx* or WT mice with Myosoft. **c** Size (CSA) distributions of *mdx* (red boxes) or WT (black circles) myofibers in aggregate (top) or by specific fiber type.

In comparing the fiber type populations between control and dystrophic muscle we discovered that the total sum of proportions of the four types exceeded 1. This is because mixed fiber types are counted as being positive for both other the subtypes contributing to the mixture (i.e. a type I/IIa fiber is counted once as both a type I fiber and a type IIa fiber). In all, there are 8 possibilities for fiber types that can be distinguished using our staining paradigm: type I, IIa, IIb, and IIx single-type fibers, and type I/IIa, I/IIb, IIa/IIb, and I/IIa/IIb mixed types. We devised a method to simultaneously evaluate the presence of all of these possibilities through extension of our original fiber typing logical array (presented in Fig. 3). Instead of assigning values of 1 to each fiber type, we chose to assign values of 1, 4, and 6 to the Types I, IIa, and IIb, respectively. With this arrangement, each of the eight possible fiber types carries a unique value (Fig. 7A). Using this method, we identified differences in mixed fiber type proportions, with WT muscle having more I/IIb and IIa/IIb mixed fibers than *mdx* muscle (Fig. 7B). CSA distributions for mixed types showed similar trends to distributions for single-type fibers: namely, *mdx* fibers had a right-skewed distribution with larger populations of very small and very large fibers, while WT fibers were approximately normally distributed (Fig. 7C).

**Figure 7.**
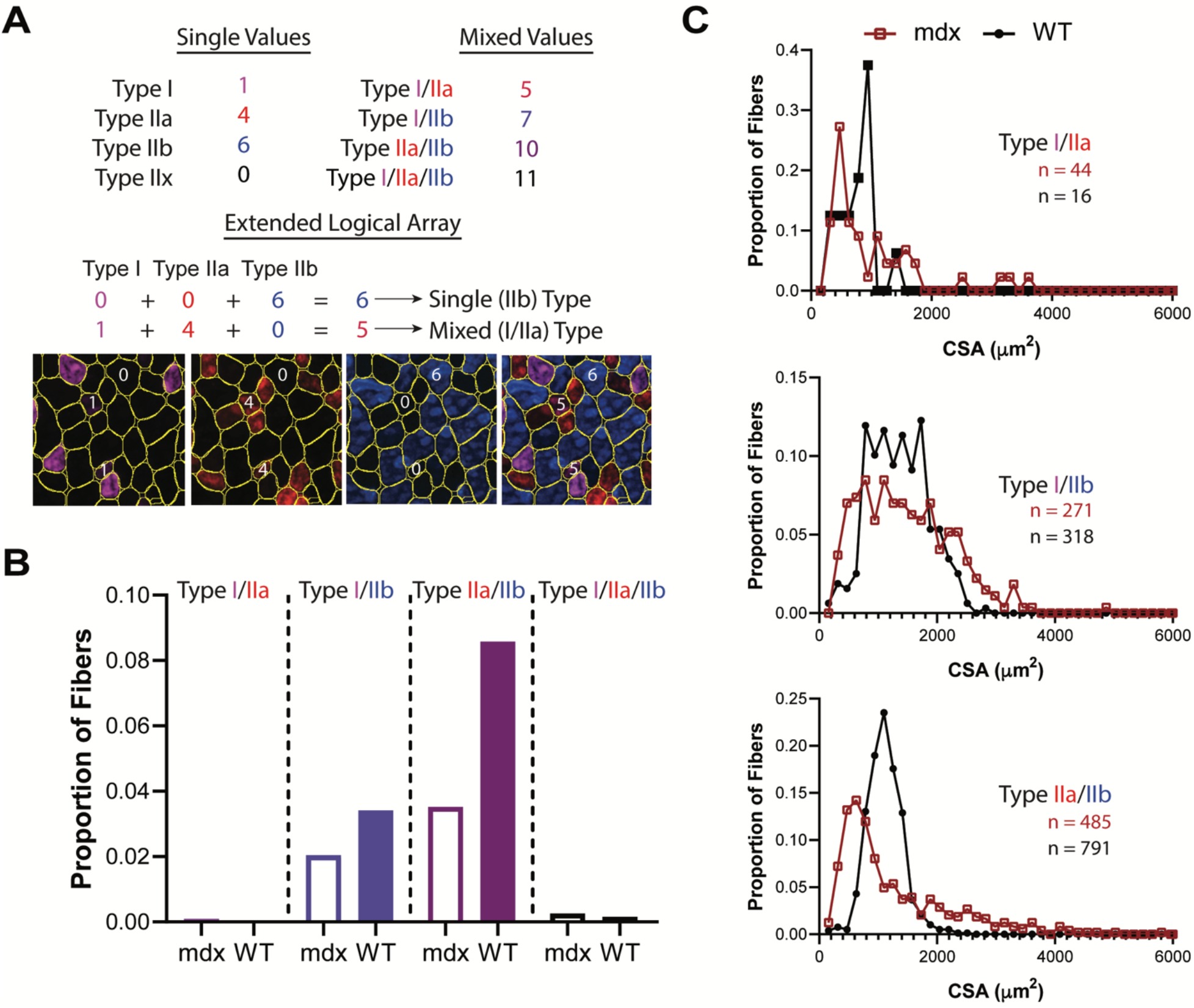
Identification of mixed fiber type populations using Myosoft. **a** The logical array used to identify Type IIx fibers (presented in Figure 3) can be extended to identify fibers of mixed type. In this case, different types are assigned unique non-zero values. When summed across channels, mixed fiber types have new, unique identifier values. **b** Mixed fiber type proportions determined by analysis of gastrocnemius/soleus sections from *mdx* or WT mice with Myosoft. **c** Comparison of CSA distributions of mixed fiber types in gastrocnemius/soleus muscles of *mdx* (red boxes) or WT (black circles) mice.

## Discussion

Histological analysis of muscle sections has been a staple of muscle physiology and neuromuscular disease research for decades; indeed, a number of neuromuscular diseases were originally named for their distinct histopathological features [25–28]. Although methods to characterize muscle fiber size and type are exceedingly common, they have been only modestly updated since their original conception. Such analyses are still routinely performed manually despite their laborious nature and consequently represent a substantial bottleneck for projects that require them. Furthermore, while manual annotation of muscle sections is not complex, there is a degree of subjectivity inherent in the measurements taken. Because of this, there can be considerable inter-individual variation which can confound the interpretation of studies reported by different research groups. Recent advances in computer technology have enabled software-based automation of standard laboratory data analysis, including analysis of digital images. To date, several groups have reported tools intended for applications in histological studies of muscle, including Myovision, SMASH, and MuscleJ [16, 18, 19]. Of these, MuscleJ has by far the most complete collection of features, with a number of capabilities that are absent even in Myosoft. However, we found that all programs tested, including MuscleJ, stumbled in analysis of large images or images with sub-optimal staining. This is emblematic of a larger problem with automated image analysis: in general, automated analysis is successful only when staining protocols are optimized to yield higher quality (greater signal:noise) input images for subsequent analysis. As a novel alternative, we employed a machine learning-based approach to improve the accuracy of image segmentation without altering standard protocols for tissue staining or image acquisition. By using two classifiers iteratively, Myosoft mathematically improves image signal:noise without the need for manipulation of original images. Altogether, our machine learning-based segmentation approach shows superior performance for accurate and efficient histology analysis compared to other muscle histology programs. In addition, Myosoft overcomes the dependence on image quality for utilizing histology analysis program.

Fiber type analysis is a common method in the muscle disease and physiology fields because it is relatively simple technically and because well-characterized antibodies are available. However, Myosoft can be used in any instance where it is necessary or desirable to make intensity measurements within identified object boundaries. In the case of muscle analysis, this might entail assessing muscle fusion *in vivo* with Pax7^cre/ERT^;tdTomato reporter mice [29], or evaluating proportions of central nuclei in a tissue section. Furthermore, while Myosoft was developed as a tool for automated analysis of skeletal muscle histology, the classifiers used in our macro could be extended to detect boundaries for non-muscle objects provided that there is sufficient distinction between the object and its boundary. Although we have not specifically tested additional applications, we posit that Myosoft could be easily modified to identify, for example, cardiomyocytes or adipocytes in histological sections.

We have shown that Myosoft yields essentially equivalent fiber size distributions to manually annotated data, but this capability is not unique. A more challenging problem in muscle histology analysis is the evaluation of specific fiber type proportions and size distributions. Identification of muscle fiber types is most commonly accomplished through immunofluorescent methods, but weak labelling or low expression of certain myosin heavy chain isoforms, as well as high background fluorescence of muscle sections at certain wavelengths, can make it difficult to obtain data that is both precise and accurate. Myosoft solves this problem by storing identified fiber boundaries as ROIs and overlaying these ROIs on images of individual fluorescent channels, each of which corresponds to a particular myosin isoform (and, by extension, fiber type). Thresholds for determining fiber types are set objectively according to the distribution of intensities for all fibers on a given channel. Although high signal:noise ratios make the task of setting the threshold simpler, we show that it is possible to determine a valid threshold even when differences between positive and negative fibers are hard to identify by eye. Furthermore, while we focus on the use of Myosoft for fiber type analysis, any standard measurement that can be made for an ROI can thus be made for a “fiber”. As an example, denervated muscle commonly features atrophic, angular fibers which could be detected by plotting distributions of morphometric features measured by Myosoft like circularity and solidity. Overall, Myosoft provides an objective platform for extracting type-specific fiber measurements rapidly and accurately.

Computing technology is now firmly integrated into the biological research enterprise, but rapid advances in fields such as machine learning and artificial intelligence offer new opportunities for automation of tedious analyses. Although Myosoft is, to our knowledge, the first program to exploit machine learning for use in muscle histological analysis, it is indeed only a first step. While the program represents a substantial improvement over manual analysis, it is not as complete as MuscleJ with respect to the kinds of myological analyses it will perform. And while MuscleJ will automatically report processed data to the user, Myosoft requires that the investigator evaluate the large numerical datasets it generates. However, it should be noted that since Myosoft provides users with raw data and reference images, it is possible to corroborate results (something that is entirely impossible within MuscleJ). Furthermore, we have attempted to alleviate the burden of data processing to some extent by providing a MATLAB script that will automatically provide single- and mixed-type proportions and areas once thresholds are set for each channel (Supplemental file 3 and 4 (tutorial)). We have likewise taken great care to ensure that Myosoft will be simple and convenient to use for the entire muscle community through extensive beta testing and by providing detailed instructions for use/troubleshooting. Looking forward, it will be interesting to extend this approach to other types of analyses, both in muscle and beyond. As the use of automation expands in biological sciences, previously intractable research questions will become increasingly accessible.

## Conclusions

Myosoft synergizes the power of machine learning-based image segmentation with thresholding-based object extraction and quantification to obtain the morphometry and type of fibers in a given histological section of muscle. In doing so, it is capable of circumventing the time, effort, and error incurred by manual histology analysis and addresses the central limitations of its peers. Myosoft is freely available in the open access image analysis platform: Fiji (Fiji Is Just ImageJ), permitting the use of the vast repertoire of functions therein which are familiar to much of the muscle community. Myosoft also applies the power and versatility of a machine learning-based approach to image analysis. We anticipate that Myosoft will be an especially useful tool for the muscle community and will serve as a scaffold for the creation of future automation programs.

## Declarations

### Author Contributions

Lucas Encarnacion-Rivera: Conception and design, code writing, collection and/or assembly of data, data analysis and interpretation, manuscript writing, final approval of manuscript

Steven Foltz: Conception and design, data analysis and interpretation, manuscript writing, final approval of manuscript

H. Criss Hartzell: Data analysis and interpretation, manuscript writing, final approval of manuscript

Hyo-Jung Choo: Conception and design, data analysis and interpretation, manuscript writing, final approval of manuscript

