## Supplemental file 2 for "Myosoft: an automated muscle histology analysis tool using machine learning algorithm utilizing FIJI/ImageJ software"

**About Myosoft**:

Myosoft is an ImageJ-based macro that can be used to analyze muscle fiber size and type almost automatically. Myosoft accepts up to 4-channel images, identifies muscle fiber boundaries and measures fiber cross-sectional area, and categorizes each fiber according to the channels available. Myosoft is freely available for download here. Myosoft is designed to be user-friendly while also retaining some ability for user input, specifically with regard to morphometric gates that the macro uses to identify objects of interest (e.g. muscle fibers). Therefore, the user will receive several prompts when the macro is run.

Myosoft is built to handle (up to) 4 channel images, with a membrane counterstain on 1 channel, and myosin heavy chain isoforms (or other intracellular proteins) on the remaining 3. By default, Myosoft will prompt the user for the channel number corresponding to the counterstain, type IIa, type I, and type IIb fibers (more detail in section 10). If you are not using all 4 channels, simply provide the numbers for the channels you are using and leave the others blank. Myosoft will still "analyze/provide results" for blank channels, but in actuality, it just duplicates analysis of channel 1, and the data for these duplicates can be ignored.

> ***Note:** the channel names are only names. They do not affect the analysis in any way, but the results will be stored in .xlsx files that bear the names of the channels. If you are not staining typeI, typeIIa, and typeIIb fibers, we would recommend running Myosoft with these default labels and simply keeping track of which of those labels corresponds to each of your real stains. It is possible to change the names within the code, but that process is slightly complicated.

A detailed description of Myosoft outputs is given in the results section below. Briefly, outputs include: (1) Images of each channel with an overlay of "fiber" boundaries (ROIs), (2) a .xlsx spreadsheet (Excel compatible) that includes all measurements taken by Myosoft for each channel, and (3) a color-coded version of the input image where fibers are pseudo-colored according to fiber cross-sectional area (CSA, a scale is also saved in the results, showing the colors assigned to fibers of various sizes).

If you would like to adapt the Myosoft code for a new application, the macro can be edited in FIJI **Plugins>Macros>Edit>Myosoft.ijm** (select the .ijm file from its source folder). The source code is annotated by the author for clarity.

**Requisites for Myosoft**

**FIJI (Fiji Is Just ImageJ) version 2.0 with Java 1.8 or newer**

- The latest version can be downloaded from https://imagej.net/Fiji/Downloads.
- The version can be checked in the Fiji application

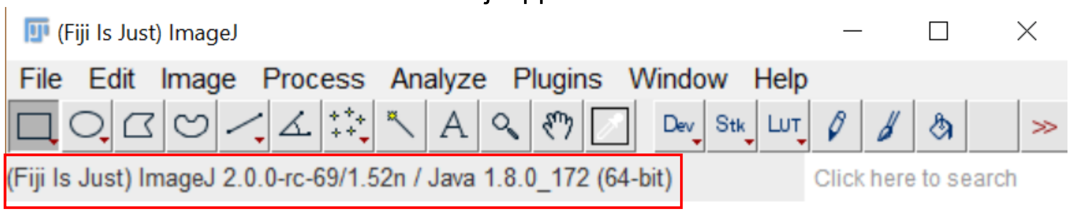


**Several FIJI plugins must be installed. These include:**

**Weka Segmentation (TWS) v3.2.33**

- This should come with the latest FIJI download and can be found in **Plugins>Segmentation> Trainable Weka Segmentation**
- Version of TWS can be found at the top left corner of the window
    - 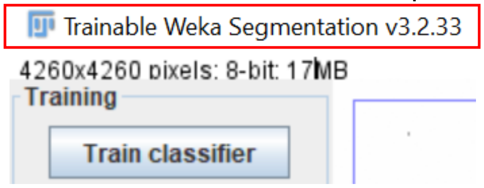


**BioVoxxel Update**

- **FIJI > HELP > UPDATE (Not "Update ImageJ")**
    - Note: when selecting "Update" from the "Help" menu, FIJI will automatically run an updater. The user must wait for the updating to complete (usually a few minutes or less) before the option to "Manage update sites" is displayed.

  **Manage Update Sites: Check BioVoxxel.**

  **Click: Add Update Site**

  **Click: Apply changes**

  **Restart FIJI.**

- This will install Watershed Irregular Features and Biovoxxel Extended Particle Analyzer

**Bar Color Coder**

- **FIJI > HELP > UPDATE (Not "Update ImageJ")**

  **Manage Update Sites: Check BAR.**

  **Click: Add Update Site**

  **Click: Apply changes**

  **Restart FIJI.**

    - Note: BAR is a suite of plugins with a variety of functionalities. Once the BAR update site is enabled, "BAR" will appear above the toolbar in the primary FIJI menu.

**<u>Using Myosoft – Windows and Mac</u>**

<u>- Download and initialization</u>

1. Myosoft is freely available for download here or from supplemental files.

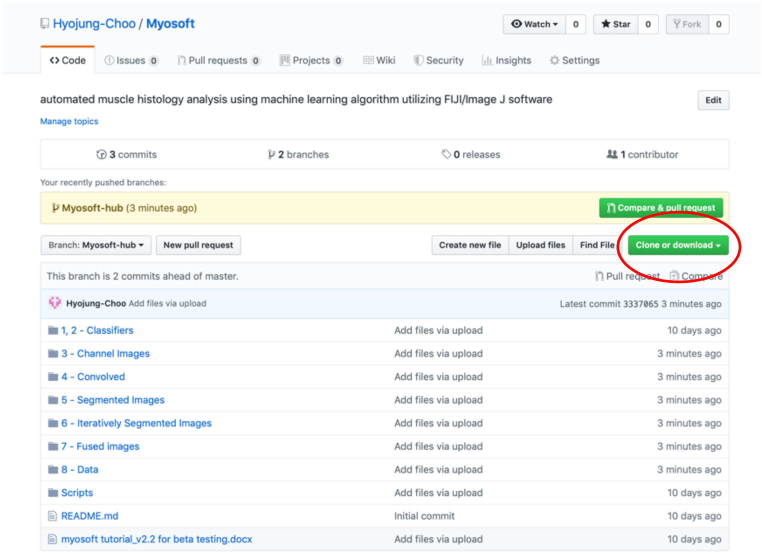


2. Save the Myosoft Hub folder to an easily accessible location
   - The directory should be (for example) C:\Myosoft Hub for Windows, Document/Myosoft Hub for Mac
   - A number of subfolders are included in the parental “Myosoft Hub” folder. These are populated when Myosoft runs.
3. Open a multichannel image in FIJI (Myosoft_Sample_Image.nd2 is in the Myosoft Hub folder). When a multi-channel image is opened in FIJI, the bio-formats Import Options window will open. Open your image as a hyperstack, and do not split channels (see red circles below). **It is highly recommended that you know the scale of your image (pixels per micron) before running Myosoft.** You will be asked to supply this value in Step 8.

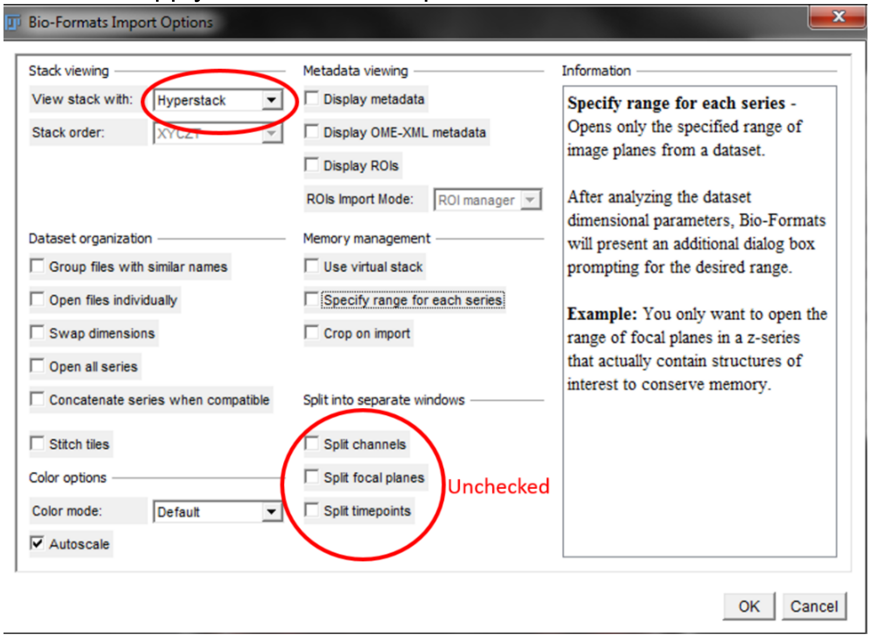

    - **If you are starting with single channel images**: We provide a macro “images to hyperstack.ijm” to combine your images and format them as a hyperstack in the Myosoft Hub\scripts folder. Save the resulting image before running Myosoft.
4. Open the Myosoft v6.ijm macro by dragging it into FIJI or by going to **Plugins>Macros>Run and navigate to Myosoft v6.ijm in Myosoft Hub\imagej scripts**.
    - Click Run.
    - First, you will receive a prompt regarding the location of the “classifiers”. The classifiers are pre-trained segmentation algorithms that Myosoft uses to detect object boundaries (this is the machine learning component of Myosoft). The default location for the classifiers is \Myosoft Hub\1,2 - Classifiers.
    - Next, you will receive several prompts regarding where to save output data. Default directories exist in Myosoft Hub, but you can make and name your own directories as you wish. Any file generated will be overwritten every time the macro is run **unless you make subdirectories within them for each image you analyze (see step 5, below)**. Prompts are:
        - “choose place to save single channel images” – Make a subdirectory in \Myosoft Hub\3 - Channel Images.
        - “choose place to save convolved images” – Make a subdirectory in \Myosoft\4 - Convolved
        - “choose place to save segmented images” – Make a subdirectory in \Myosoft Hub\5 - Segmented Images
        - “choose place to save iteratively segmented images” – Make a subdirectory in \Myosoft Hub\6- Iteratively Segmented Images
        - “choose place to save fused images” – Make a subdirectory in \Myosoft Hub\7 - Fused Images
        - “choose place to save fiber type and morphometry data” – Make a subdirectory in \Myosoft Hub\8 – Data

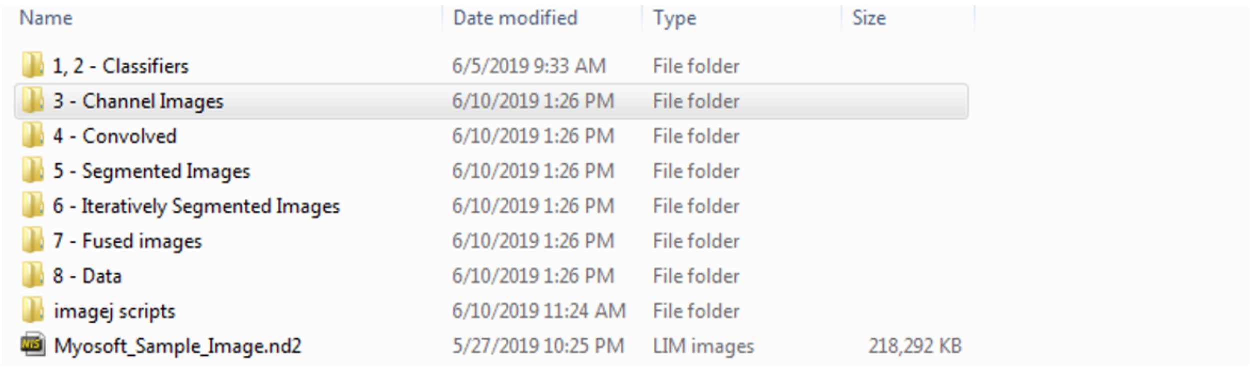


**Above**: Default Myosoft directories.

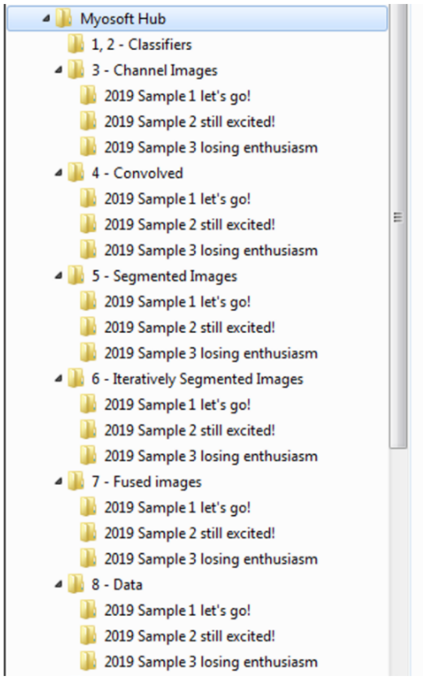


**Above**: Example subdirectories for multiple samples.

5. Morphometric gates

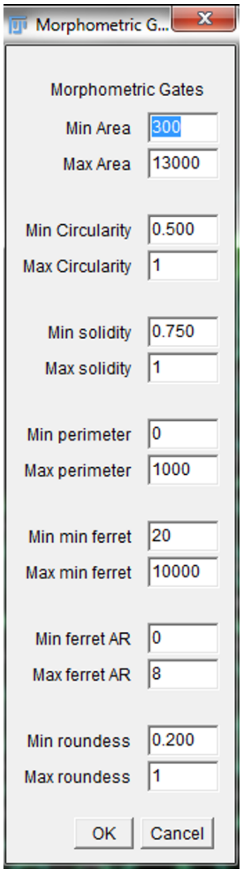


**Above:** Default morphometric gate values

- Myosoft uses the Extended Particle Analyzer in the BioVoxxel Toolbox to set a number of "gates," or exclusion criteria for objects that are classified as myofibers but are probably not myofibers or *are* myofibers, but in an improper orientation. For example, the minimum area gate would exclude small blood vessels or nerves, while the maximum area and circularity gates would exclude oblique or longitudinal myofibers within the section.
  a. Min/Max area – default range is 300-13000μm$^2$. Ideally, this range should be tested on a control section, and small and large values should be examined to ensure that these gates are appropriate for your sample. As a general rule, fibers of ~5000μm$^2$ are quite large and appear at the upper end of ranges reported in the literature. (Perimeter)$^2$
  b. Min/Max circularity – Circularity = $\frac{4\pi(Area)}{(Perimeter)^2}$; a two-dimensional area:surface measurement. For a perfect circle, this value would be 1.
  c. Min/Max solidity – Solidity = $\frac{Area}{Convex\ Area}$; in other words, what is the ratio of the true area of your object to the area it would have if it were imagined as a convex polygon (concavities extended outward)?
  d. Min/Max perimeter – measures the length of the object perimeter.
  e. Min/max minimum feret distance – minimum feret distance is the shortest possible distance between any two tangent lines of your object perimeter. For an ovoid shape, this would be the length of the short axis.
  f. Min/max feret AR – the aspect ratio (AR) of ferets for your object, that is, $\frac{FeretMax}{FeretMin}$. For an ovoid shape, this would be the ratio of the axes.
  g. Min/max roundness – Roundness = $\frac{4Area}{\pi(\max diameter)^2}$; this formula compares how closely your area matches the area of a perfect circle (value for perfect circle is 1).
- The Extended Particle Analyzer allows for more gates than those included as defaults in the Myosoft code. To view these, and to see some pictoral representations of the gates used, check here.

6. A prompt will appear asking for an ROI expansion factor. See Fig. 1, Step 8 in the paper to see an example of 4 pixel ROI expansion. Any integer value (positive or negative) is acceptable (negative values will shrink ROIs). We recommend starting with a value of 4, which is the default, and adjusting only if this does not suit your needs.
   - Why is this done? The binary, segmented images show what the classifier views as the probability that a given pixel in the image represents the boundary of an object (black p=1, white p=0). Typically, the classifiers will view several pixels at the cell boundary as plausibly marking the boundary, which is why the iteratively segmented images appear to have thick black borders around the fibers. ROIs denoting the fiber edges are drawn at the innermost edge of the probability map, which is usually well within the cell. ROI expansion moves the ROI outward in all directions by the number of pixels chosen with this prompt.

7. A prompt will appear asking you for the scale of your image. **IMPORTANT: you must know the scale (pixels per micron) of your image before running Myosoft.** If you do not supply the correct scale, then CSA values generated in Myosoft will not be in real spatial units. As a default, a value of 0.9091px/micron is provided (**this is the scale value in the sample image provided in \Myosoft Hub**), but it is likely that your image is scaled differently.
8. A prompt will appear asking you for Color Coder parameters.
    - Once Myosoft has completed fiber CSA analysis, it will automatically run an ROI Color Coder plugin (from BAR suite of plugins). This will provide a color-coded mask of your input image where fibers are assigned colors based on their CSA. This is an aid for visual inspection of your sample's fiber CSA distribution, which is meant to complement a graphical representation of CSA values.

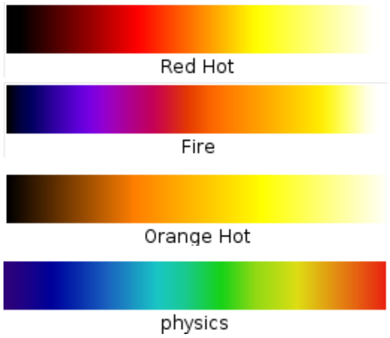


Small -----------------------> Large
**Above:** Color code LUT options.

        a. Minimum value – the lowest area value to which you want to assign a unique color. All objects below this value will receive the same color as the minimum.
        b. Opacity – opacity value for colorization. The color-coded ROIs are overlaid on your original image (but saved under a different name), so if you wish to be able to see the original image objects, use a lower opacity value (that is, less than 50). Normally, this is unnecessary, and a value of 80-100 will nicely mask your image.
        c. Maximum Value – the highest area value to which you want to assign a unique color. All objects larger than this value will receive the same color as the maximum. Default is 4000
        d. Color code – this is the LUT that will be applied to the data. The options are: Red Hot, Fire, Orange Hot, and physics.
    - The color-coded mask is saved in the location you chose for data (step 3 above) as "InputImageNameHere_referenceImg_ColorCode".
9. A prompt will appear asking for the position (within the hyperstack) of the counterstain channel, the type IIa, type I, and type IIb channels. 1 corresponds to the left-most position, and 4 corresponds to the right-most position. You can check the position using cursor keys or the scroll bar at the bottom of hyperstack image window.
10. The input image will next be cut in 16 sub-images. This step is included to guarantee that your machine has the necessary computing power for Myosoft to operate. Running large images through the classifier is resource intensive and inefficient. Instead, Myosoft will run 16 smaller images through the classifier consecutively. If you want to change this you will need to add/or substract lines in the code.
    - A prompt will appear asking how many divisions you would like to make of your input image. The default of 4 will cut the input image into 16 (4^2) subimages. However, Myosoft will accept any value from 2 to 5 (resulting in 4-25 sub-images). Choose higher numbers (4 or 5) if the starting image

is large and your computer has ≥8GB RAM. If the starting image is small, fewer divisions are necessary.

11. When the macro is finished, a pop-up box will display the message "Analysis Complete." Analysis of ~10000 fibers (starting image ~6x6mm) takes 30 minutes on a computer with an i7 (3 GHz) processor and 16 GB RAM.

<u>- Results</u>

12. Myosoft will generate individual spreadsheets (.xlsx format) with data from each channel in the original input image.
    - The file from the membrane counterstain will be called "Fiber_Area_Results."
    - Each of the other files will be named with the stained myosin heavy chain isoform in the title (e.g. "Fiber_type_Results_typeI").
    - In these files, each row represents a fiber identified by Myosoft, and all columns represent data for that fiber. Mean, StDev, Mode, Min and Max refer to the measured intensity from within the bounds of the fiber, while most of the other values are physical parameters (e.g. area, x/y position, min. feret). **The number in column A is the object index and is constant across all the files.** This means that object 1 in "Fiber_type_Results_typeI" is the same as object 1 in "Fiber_type_Results_TypeIIa". Further, it is the same as the object 1 circled in all reference images.
13. Myosoft will generate a series of reference images. The reference images are single channel images from the original input image with "fiber boundaries" and numbers overlaid as a mask. The number in each object corresponds to the same number in the data spreadsheets. These images are provided so that users can visualy inspect the objects from which measurements were taken by Myosoft.
14. Myosoft will generate a color-coded version of your input image where different colors indicate different sized fibers. A scale bar is also generated and saved in the location the user specifies for data (Step 6f).
    - You may choose from 1 of 4 LUTs (see 9d, above).
    - A default range of 0-4000 is set for fiber sizes. This means that fibers of ~no size will appear black, while fibers 4000um or greater will appear white. You may wish to adjust this range to better fit your own data.
15. Myosoft returns one .xlsx file for each image channel, one reference image for each channel (with ROI mask overlay), and a color-coded version of the image.

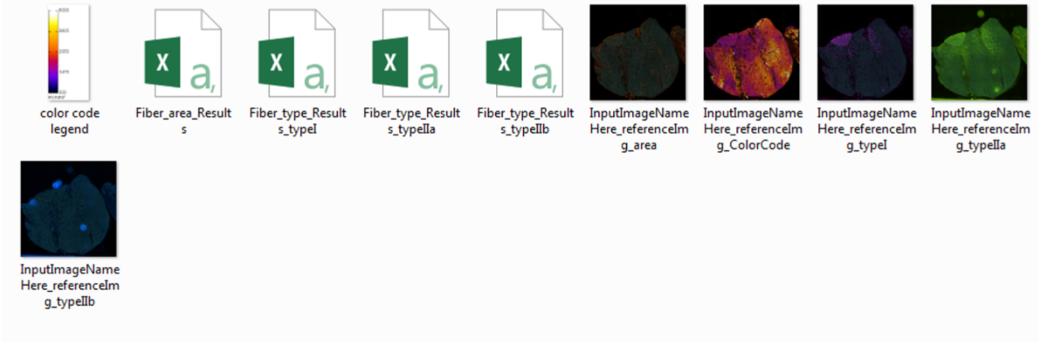


**Above**: Data folder contents.

### About the Author

Lucas Encarnacion-Rivera is currently an undergraduate student at Emory University. He may answer questions about Myosoft or help with troubleshooting if his schedule allows.
