## Supplemental file 4 for "Myosoft: an automated muscle histology analysis tool using machine learning algorithm utilizing FIJI/ImageJ software"

### Using MATLAB to analyze data generated by Myosoft

#### Introduction

Myosoft is an image analysis macro executed in ImageJ. Its primary use is for automation of skeletal muscle fiber size and type analysis. However, while Myosoft generates the pertinent data for this purpose, it is still incumbent upon the user to process the data. **fiber_processing.m** is a MATLAB script that will autonomously perform most of necessary data processing.

#### Getting started

1. Open the fiber_processing.m file in MATLAB.
2. Import Myosoft-generated data
    a. On the left side of the MATLAB window, you will see a directory. Use this to go to the folder containing Myosoft-generated data.

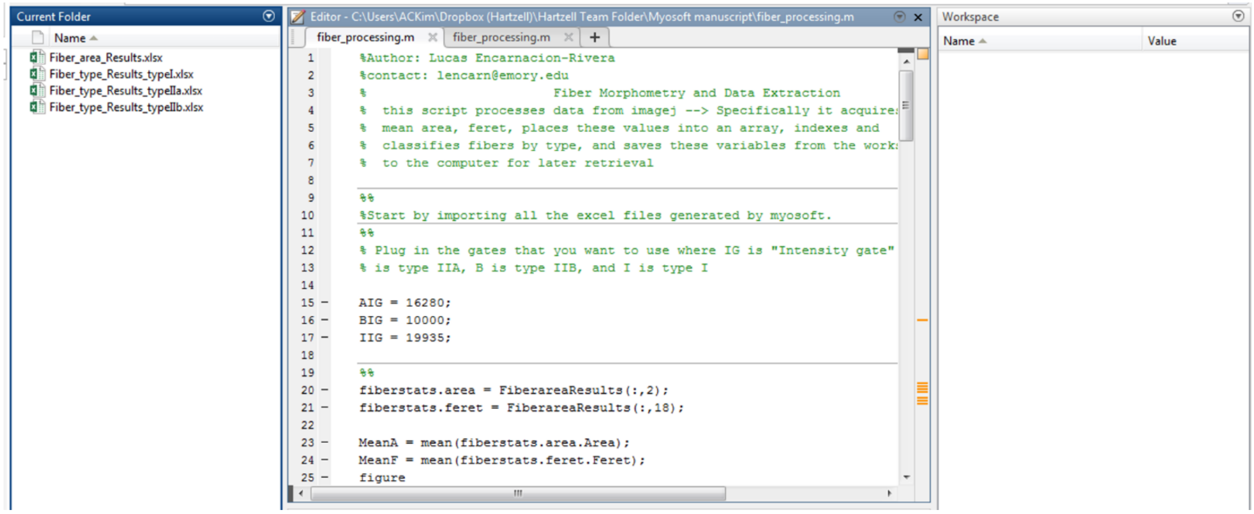


    b. Select all of the .xlsx files in this folder (there should be 4) using ctrl+click. **DO NOT CHANGE THE NAMES OF THESE FILES FROM THE DEFAULT.** They should be "Fiber_area_Results", "Fiber_type_Results_typeI", "Fiber_type_Results_typeIIa", and "Fiber_type_Results_typeIIb".
    c. Right-click on the highlighted files and choose "Import Data". A new window will pop up displaying the contents of the chosen files. In the top right of this window is a button to

"Import Selection". Choose this option for each of the 4 datasets. When you are finished, you can close out of this window.

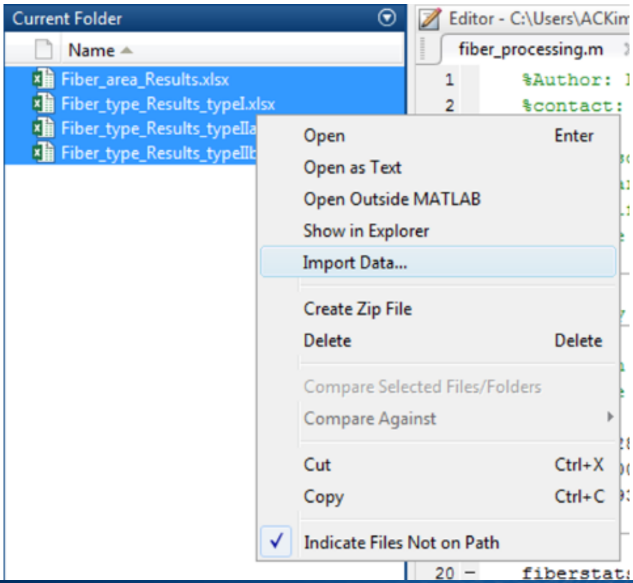


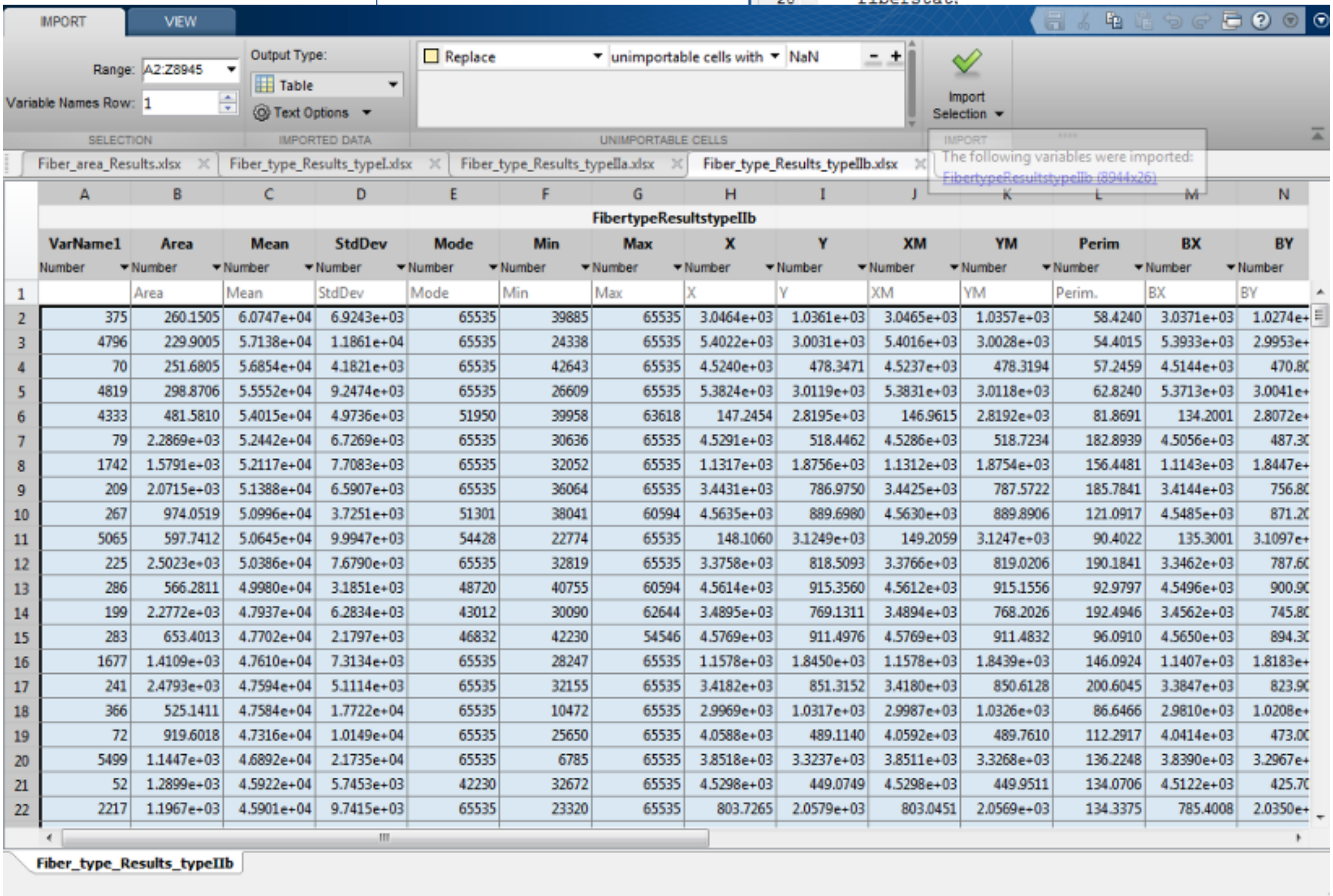

d. In the original MATLAB window, there is a "Workspace" on the right side. This should now be populated with the data you just imported.

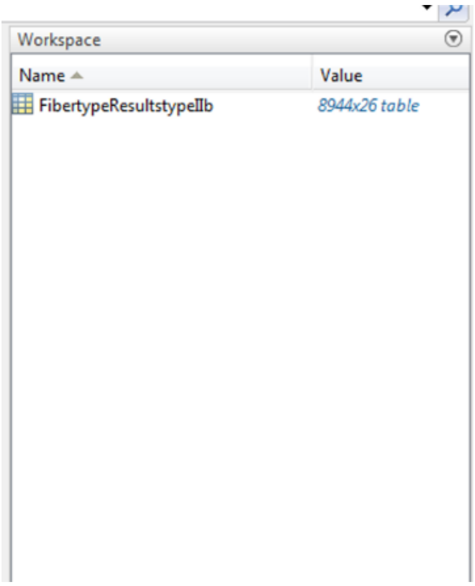


**Processing the data**

3. Click Run. Myosoft will take mean intensity measurements on all image channels. When all intensity values for a given channel are plotted as histograms, they typically display a bimodal distribution with peaks for background (low intensity) and stain-positive (high intensity) objects. Frequency plots of intensity values for each fiber type are generated in MATLAB to help you determine the intensity values you wish to use to distinguish background and stain-positive objects (i.e. the "intensity gate"). A breakpoint is set in the script after generating the histograms so that you have time to inspect them and choose thresholds.

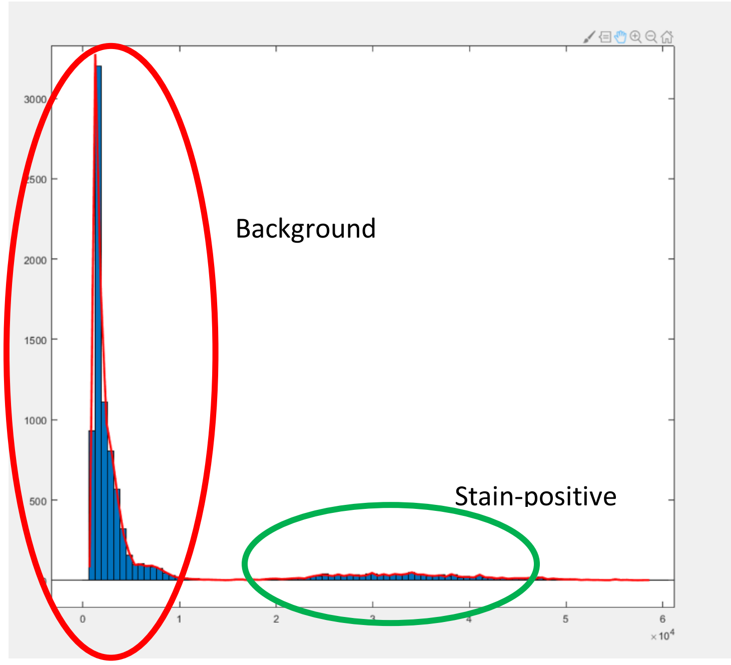

4. Set the intensity gates. Click continue.
    a. The first lines of code will read “AIG = ….” “BIG = ….” And “IIG = ….”. These stand for the intensity gate to be applied for type IIa, type IIb, and type I fibers, respectively (so AIG means **A** **I**ntensity **G**ate, referring to type II*A*).

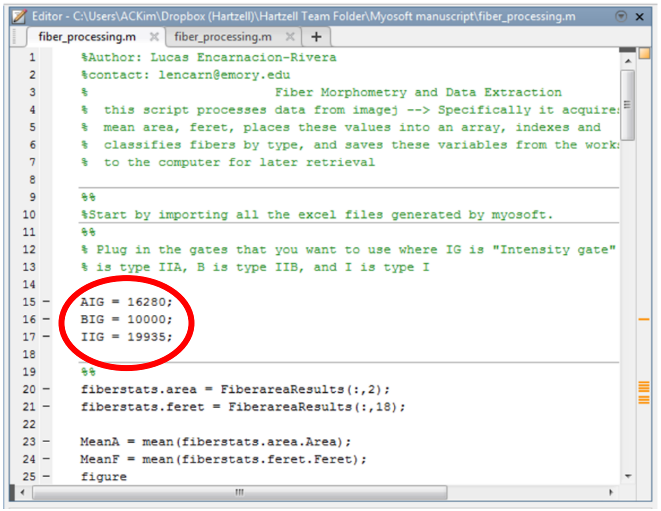


Depending on the bit-depth of the images you used in Myosoft, the range of values should be 0-256 (8-bit) or 0-65536 (16-bit) (ImageJ may open your 12-bit images as 16-bit).

    b. fiber_processing.m will use the numeric values entered to determine which objects do and do not correspond to different fiber types.

**Results and further analysis**

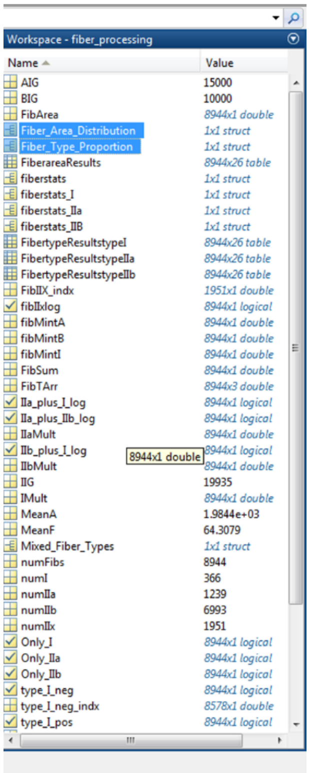


5. The workspace on the right should now be populated with several new fields. Some of these contain relevant processed data. There are 2 fields of interest.
    a. **Fiber_Area_Distribution (1x1 structure)**: This structure will contain a spreadsheets for every fiber type (single or mixed) with a single column corresponding to the CSA values for fibers of that type.
    b. **Fiber_Type_Proportion (1x1 structure)**: A simple list of the calculated proportions of Type I, IIa, IIb, and IIx fibers.

    Other fields in the workspace are used by MATLAB to obtain the processed data, but won’t be particularly useful to the user by themselves.
